# Reduced frequency of clonal hematopoiesis in Parkinson’s disease patients

**DOI:** 10.1101/2023.05.04.539397

**Authors:** Shixiang Sun, Daisy Sproviero, César Payán-Gómez, Zhenqiu Huang, Jan H.J. Hoeijmakers, Alexander Y. Maslov, Pier G. Mastroberardino, Jan Vijg

## Abstract

To test if clonal hematopoiesis of indeterminate potential (CHIP) is associated with the incidence of Parkinson’s Disease (PD), we analyzed blood whole exome sequencing data from 171 healthy controls and 335 PD subjects in the Parkinson’s Progression Markers Initiative. We observed an age-related increase of CHIP carriers in the healthy controls. Surprisingly, the percentage of CHIP carriers was significantly lower in old PD patients than in age-matched controls.

## Main

Parkinson’s disease (PD) is the second most common neurodegenerative disorder, affecting 2% of the population over 60 years old (1). The major neuropathological hallmarks of PD are the loss of dopaminergic neurons in the substantia nigra, and the aggregation of Lewy bodies containing α-synuclein. Evidence has been obtained that the molecular basis of PD involves mitochondrial dysfunction leading to an increased production of reactive oxygen species and oxidative DNA damage. This could increase somatic mutation frequency during aging and clonally amplified somatic mutations have indeed recently been reported for different regions of the brain, affecting genes enriched in synaptic and neuronal functions as well as in blood of PD patients (2).

In blood, clonal expansion of somatic mutations in hematopoietic stem cells, known as Clonal Hematopoiesis of Indeterminate Potential (CHIP), increases with age and has been associated with cancer, cardiovascular disease and mortality in general (3). CHIP clones often carrying cancer-associated somatic mutations are also linked with non-malignant diseases of aging (4), including neurodegenerative disease (5). However, in Alzheimer’s disease (AD) patients, CHIP frequency has been found to be lower, not higher, than in age-matched controls, with CHIP carriers showing reduced risk of dementia or neuropathologic features (6). Here, we hypothesized that the incidence of CHIP in PD patients would also be lower than in healthy, age-matched controls.

We assessed CHIP frequencies in whole exome (WES) data from blood of PD patients and control subjects available in the Parkinson’s Progression Markers Initiative (PPMI; https://www.ppmi-info.org) (**Fig. 1A**). In this cohort, there is no significant difference in age between PD patients and healthy controls, with the ages at enrollment ranging from 40 to 79 years (P = 0.253; **Table S1**). Sex distributions are consistent between PD patients and healthy controls (P = 0.921; **Table S1**). Only PD patients with demonstrated dopaminergic deficits were selected for this study.

**Fig. 1.**
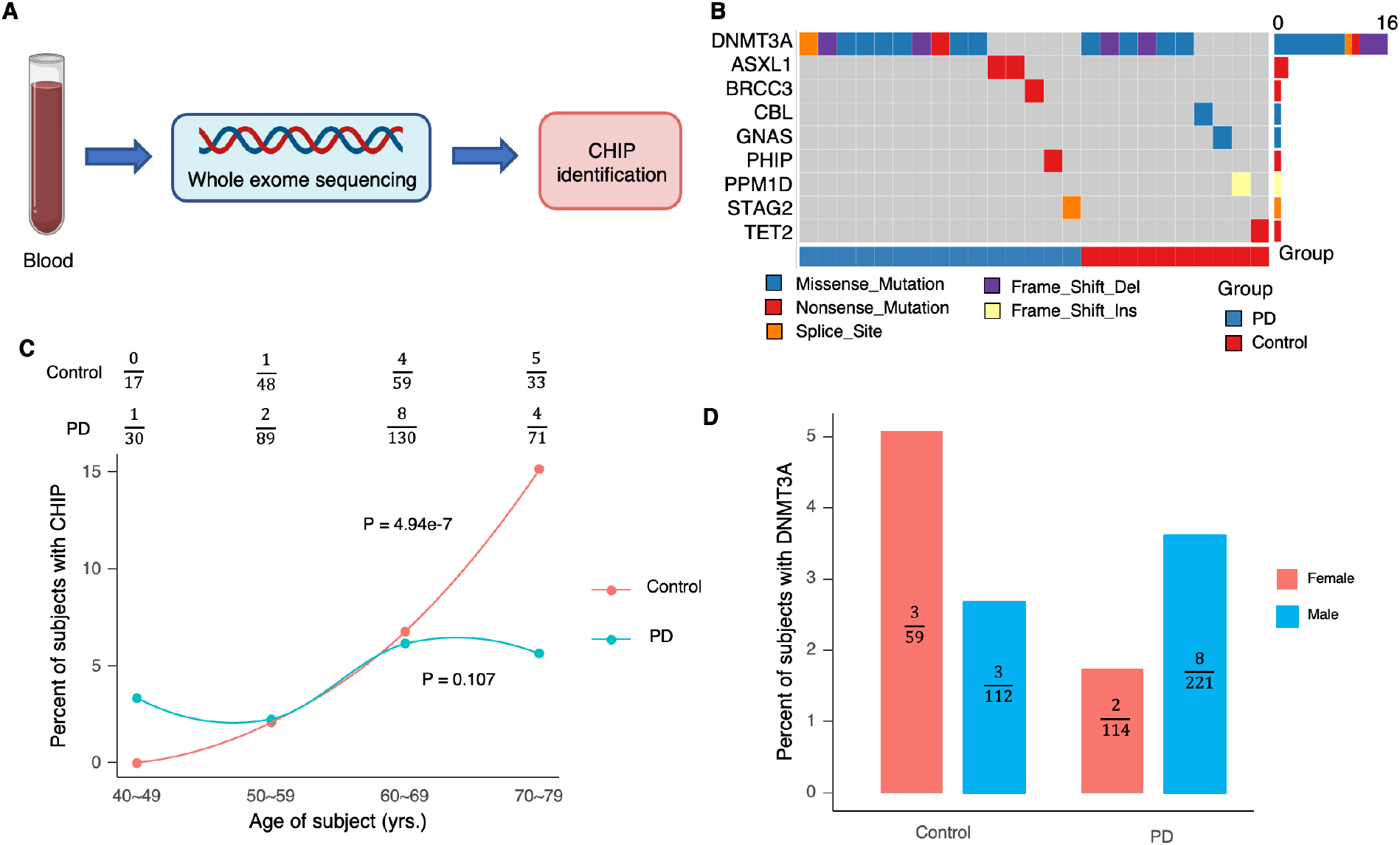
Study overview and CHIP variants identified in WES data. (A) Workflow of sequencing and data processing. Sequencing data was obtained from PPMI. The figure was created using BioRender. (B) Co-mutation plot with mutations across the 25 subjects from healthy controls and PD patients. (C) Distribution of percentages of CHIP carriers across different age bins. The numbers in the top panel indicated the numbers of CHIP carriers and total subjects in the age bin for healthy controls and PD patients, separately. (D) Percentage of DNMT3A mutation carriers in female and male subjects of two cohorts.

We examined known CHIP variants in 73 genes in the WES data involving 171 healthy controls and 335 PD subjects (Supplementary Methods). The sequencing depth was 47.6±20.3 and 51.0±18.6 in healthy controls and PD patients, respectively. A variant was considered a CHIP variant if its allele frequency was larger than 2% and the variant was supported by at least 7 reads (Supplementary Methods). Of the 506 subjects, we identified 25 CHIP events in 25 subjects (**Table S2**). The top affected gene was DNA methyltransferase 3A (*DNMT3A*; **Fig. 1B**), which is the major affected gene that has been associated with CHIP (3).

Then, we examined if the percentage of CHIP carriers was associated with age or PD status. As only 5% of the subjects were CHIP carriers, we summarized the results by age bins (**Fig. 1C**). The results indicate that the percentages of CHIP carriers are significantly increased with age in controls (P = 4.94e-7) but not in PD samples (P = 0.107), confirming the well-documented age-related increase of CHIP in blood (3). Further, we found that the relationship between age and the percentages of CHIP carriers are significantly different between the healthy controls and PD patients (P = 1.06e-3). In other words, as age increases, the likelihood of acquiring CHIP variants is lower in PD patients compared to healthy controls, leading to a lower percentage of CHIP carriers among the oldest age group of PD patients (Ratio = 0.37, P = 0.020).

Next, we compared the most highly mutated CHIP gene between female and male subjects from controls and PD patients, separately (**Fig. 1D**). In the control group, we observed a trend of a higher percentage of DNMT3A carriers in females than in males (Ratio = 1.94, P = 0.384). Conversely, in PD patients, the percentage of DNMT3A carriers was lower in females than in males (Ratio = 0.48, P = 0.063). These reversed trends suggest a different mutational pattern of genes in blood from PD patients compared to healthy controls.

In summary, our results indicate that the percentage of subjects with CHIP is increased in blood in controls, which is well-documented (3). However, in contrast to previously observed associations of CHIP levels with risk of age-related, chronic disease, most notably cardiovascular disease and overall mortality (4), the percentage of CHIP carriers was found not to be increased in the oldest PD patients compared with oldest controls. These results suggest that either PD protects against CHIP or CHIP protects against PD. This observation is in keeping with recent findings of CHIP in Alzheimer’s disease (AD), indicating that the frequency of CHIP carriers in patients is lower, rather than higher, compared to age-matched control subjects (6). Additionally, CHIP has recently been reported to have a negative association with diabetic peripheral neuropathy in Type 2 Diabetes (7). It should also be noted that PD has previously been found negatively associated with multiple cancers, including leukemia (8). Thus, it is possible that individuals without CHIP have a lower cancer risk and a higher likelihood of aging and developing PD.

## Methods

### Data source

Basic information of subjects was obtained from PPMI (file name: meta_data.11192021.csv; https://ida.loni.usc.edu/web/ppmi-rnaseq-app#Accessdata). Missed age and sex information was found in (9). Raw exome sequencing data was obtained from PPMI’s data server.

### Examining CHIP variants

Raw sequencing data were trimmed for adapter and low-quality nucleotides with Trim Galore (version 0.6.7). The minimal length of reads for alignment was required to be at least 30 bp. Alignments were performed according to the best practices of the Genome Analysis Toolkit (GATK version: 4.2.5.0) (10). In general, trimmed reads were aligned with BWA (mem mode; version 0.7.17) (11). Removal of PCR duplications and base recalibration were performed in the alignment files using GATK. Mutect2 (version 4.2.5.0) (12) was applied to call known variants in known CHIP associated genes obtained from (13). Candidates were kept with pre-applied filters (14): if they passed Mutect2 default filtering, and had a read depth ≥20x, the number of reads supporting variant alleles ≥7 in single nucleotide variants and ≥10 in other mutation types, the variant allele fraction (VAF) ≥2%, gnomAD allele frequency <0.001, and at least one read in both forward and reverse direction supporting the reference and variant alleles.

### Sex as a Biological Variable

The study includes both male and female subjects sequenced by the PPMI. The sex distribution is the similar between healthy control (Male: 65.5%) and PD patients (Male: 66.0%). Our primary focus is not the influence of sex on the emergence of the CHIP mutation, and the analysis is conducted on age group level. Hence, sex was excluded as the factor in the statistical model to simplify the analysis and focus on age-related differences. Additionally, statistical analysis was limited to driver genes specifically, rather than including cohort-level factors.

### Statistical analyses

To compare the distribution of age and sex between PD and healthy controls, two-tailed student’s t-test and Fisher’s exact test were used, correspondingly. A generalized linear regression model was applied to examine the effect of age or PD status, including intersection of PD status on age increase, with Likelihood Ratio Test to assess the statistical significance. Exact Binomial test was used to compare the percentage of CHIP carriers in oldest groups of healthy control and PD subjects, and applied to the percentage of DNMT3A carriers between female and male groups in healthy control and PD patients.

## Data availability

All raw sequencing data were obtained from https://www.ppmi-info.org. Additional data and script related to this paper may be requested from the authors.

## Acknowledgments

We thank all members of the Jan Vijg’s laboratory for helpful discussions related to this project. We thank the staffs in the PPMI for assisting the data access.

Data used in the preparation of this article were obtained on [2023-02-23] from the Parkinson’s Progression Markers Initiative (PPMI) database (https://www.ppmi-info.org/accessdata-specimens/download-data), RRID:SCR_006431. For up-to-date information on the study, visit http://www.ppmi-info.org.

PPMI – a public-private partnership – is funded by the Michael J. Fox Foundation for Parkinson’s Research and funding partners, including 4D Pharma, Abbvie, AcureX, Allergan, Amathus Therapeutics, Aligning Science Across Parkinson’s, AskBio, Avid Radiopharmaceuticals, BIAL, BioArctic, Biogen, Biohaven, BioLegend, BlueRock Therapeutics, Bristol-Myers Squibb, Calico Labs, Capsida Biotherapeutics, Celgene, Cerevel Therapeutics, Coave Therapeutics, DaCapo Brainscience, Denali, Edmond J. Safra Foundation, Eli Lilly, Gain Therapeutics, GE HealthCare, Genentech, GSK, Golub Capital, Handl Therapeutics, Insitro, Jazz Pharmaceuticals, Johnson & Johnson Innovative Medicine, Lundbeck, Merck, Meso Scale Discovery, Mission Therapeutics, Neurocrine Biosciences, Neuron23, Neuropore, Pfizer, Piramal, Prevail Therapeutics, Roche, Sanofi, Servier, Sun Pharma Advanced Research Company, Takeda, Teva, UCB, Vanqua Bio, Verily, Voyager Therapeutics, the Weston Family Foundation and Yumanity Therapeutics.

## Author contributions

J.V. and P.G.M. conceived and supervised the study. S.S. analyzed the data. S.S., D.S., C.P.G., Z.H., J.H.H, A.Y.M, P.G.M., and J.V. wrote the manuscript.

